# Estimation of stature from hand length and hand breadth in undergraduate medical students: An anthropometric study

**DOI:** 10.1101/2025.06.03.657765

**Authors:** Niraj Pandey, Navindra Phuyal, Anusuya Shrestha, Anup Pandeya, Ananda Kumar Mishra, Kumar Bhushal

## Abstract

**Background:** Stature, a determinant that aids in personal identification, can be estimated using measurements of various body parts. This study measured the length and breadth of hands and the height of the participants, aiming to develop linear regression equations for stature estimation.

**Methods:** This is a cross-sectional descriptive study in which 195 medical students aged 18 to 24 were enrolled, and measurement of stature was done by stadiometer and hand length and breadth by vernier calliper, after an informed written consent was obtained.

**Results:** Mean stature was higher in males (170.24±5.79 cm) as compared to females (159.01±7.261 cm). Mean length and breadth of both right and left hands were also found to be higher in males when compared with females. With a standard error of estimates of 5.083 cm and an R^2^ of 0.648, the multiple linear regression analysis’s model summary concluded that R is .805, indicating a strong positive association between stature estimation and hand dimensions.

**Conclusions:** Hand length and hand breadth together may be a powerful predictor of stature, as indicated by the greater R value when compared to regression coefficients derived for other body parts. A variable entity, height is influenced by genetics and nutritional status. Future studies can be done taking these factors into consideration and also including a larger population size.

## 1. Introduction

Age, sex, stature and ethnicity are considered the “big four” for personal identification.[1] The natural height of a person who is standing upright is taken as their stature, and it is determined by a variety of factors, like hereditary and environmental.[2] Stature is a feature valuable for ‘tentative identification’ and also an important parameter for medico-legal examination and deduction of the identity of the individual.[3–9] Stature can be used in verification of the identity of individuals in cases like disaster victims.[4] Forensic scientists often find dismembered body parts in trenches, rubbish and elsewhere that are brought for examination, leading to challenges in identification. [4,10] Estimation of stature could be really helpful in those conditions too.

Many statural tables have been formulated that help anthropologists and forensic experts to estimate stature.[11] Additionally, measurements of various body parts and proportions are being used and, among all long bones are the most prevalent.[7] A comparison of the relationships between body segments is used by anthropologists to indicate variations among the origins of various ethnic groups. It is also valuable in finding the association between various lifestyles, usage of energy and locomotion patterns.[12] Although there are established relationships between body parameters, they tend to vary between populations of different regions or different ethnicities that are affected by the level of physical activity and the status of nutrition.[12,13] Stature estimation has been done using foot length in this region that showed a significant correlation.[14] Hand length alone was employed in another investigation to estimate stature while hand breadth was not taken into account.[15] A similar study was done by Shrestha S et al. where difference in right and left hands was not addressed.[16] Regression equations may be affected by differences in measurements of right and left sides. In this study, both hand length and breadth were included and differences in right and left sides were also considered that eventually derived a more robust regression equation and address the research gaps.

This study measured the length and breadth of hands and stature of the individuals. It aimed to estimate the stature from dimensions of the hands and see if there is any correlation between them. It also aimed to find if there are differences between males and females regarding the dimensions of hand, stature and its estimation.

## 2. Methods

### 2.1 Sample size and consent

A cross-sectional, observational study was carried out. It was done on medical students of Devdaha Medical College and Research Institute from 3^rd^ March 2024 to 14^th^ April 2024, after ethical approval was taken from the Institutional Review Committee of Devdaha Medical College and Research Institute, Devdaha, Rupandehi, Nepal, with approval number: 02/2024 (Ref No: 94610801082).

The sample size was calculated using the formula:

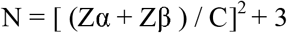

Where:

- Zα=1.96 (95% confidence interval)
- Zβ=0.84 (80% power)
- C=0.5 ln{(1+r)/(1−r)}
- r = 0.34 (Coefficient correlation) [3]

The minimum sample size for each sex was found to be 66. This study, included 195 students (80 female and 115 male). Participants were explained about the objectives of the study and the procedure of data collection and also about privacy and confidentiality. They were also informed about their right to refuse and withdraw consent at any time. The measurements were taken only after voluntary written informed consents were obtained.

Participants with deformities of the hand and vertebra, with a history of trauma to the hand and with features of dysmorphic syndromes were excluded. All other students who consented were included in the study.

#### 2.2 Data collection technique

Stature was measured using a stadiometer to the nearest centimetre (cm). Study participants were asked to stand erect in anatomical position, barefoot. The head was oriented in Frankfurt’s plane, and the measurement was taken from the vertex of the head to the sole of the foot.

Participants were asked to rest their hand against a flat cardboard in a supine position with fully extended fingers. The thumb was kept in extension while all other fingers were adducted. The length of the hand was taken from the inter-stylion point, i.e., the mid-point of the line joining the most distal point of the radial styloid process (stylion radiale) and the most distal point of the ulnar styloid process (stylion ulnare), to the most distal tip of the third finger. Breadth of the hand was taken from the most medially projected end present on the head of the second metacarpal with the hand stretched, i.e., metacarpal-radiale, to the most projected point laterally on the head of the fifth metacarpal with the hands stretched, i.e., metacarpal-ulnare, as shown in Fig 1. Measurements were taken using a vernier calliper.

**Fig 1:**
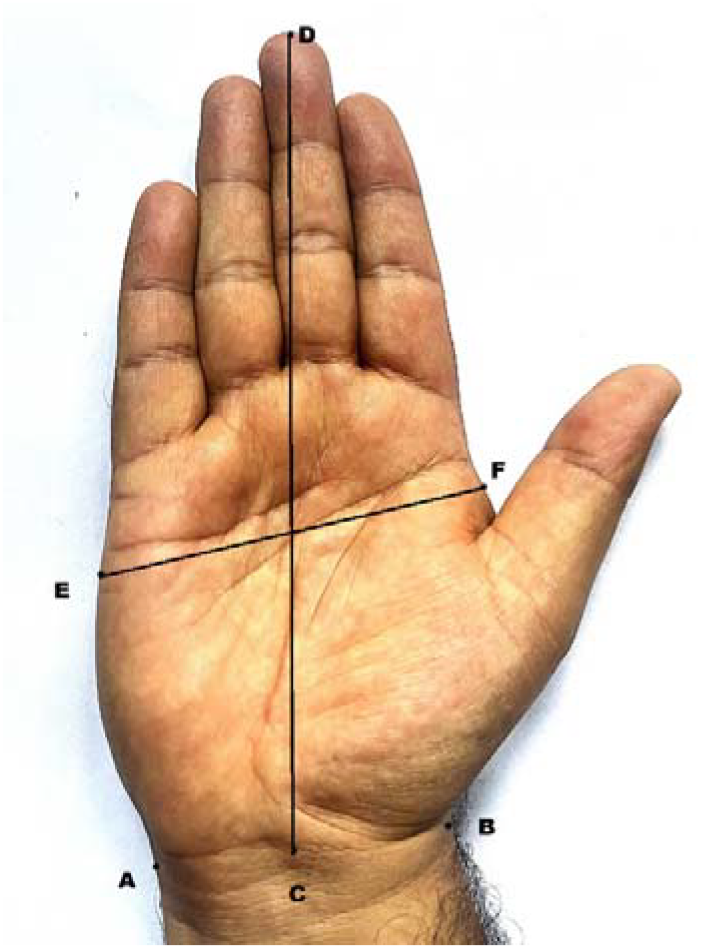
Measurements of hand. A = stylion ulnare, B = stylion radiale, C = Inter-stylion, D = tip of third finger, E = metacarpal radiale, F = metacarpal ulnare, DC = Length of hand, EF = Breadth of hand.

### 2.3 Data reliability

All the measurements were taken three times by a single well-versed investigator, and an average was calculated and noted in the pro forma. Height measurements were made at noon, taking into account variations in height during the day. In order to evaluate intra-observer reliability, in a subset of 22 participants measurements were taken again two days later, and TEM, rTEM, R and ICC were calculated for each parameter, as displayed in **Table 1**. The results show that the R value range from 0.94 to 0.98, indicating highly reliable data, and the ICC was ranging from 0.89 to 0.97, displaying consistency in the data. Data entry was done in a Microsoft Excel spreadsheet and later transferred to SPSS.

**Table 1:** Intra observer TEM, rTEM, Reliability Coefficient and Intra class correlation coefficient.

| Parameters | T | rTEM | R | ICC |
| --- | --- | --- | --- | --- |
| LHB | 0.108711 | 1.372147 | 0.976824 | 0.964635 |
| RHB | 0.107661 | 1.337027 | 0.981709 | 0.96818 |
| RHL | 0.173205 | 0.977556 | 0.984673 | 0.970136 |
| LHL | 0.338446 | 1.89944 | 0.941831 | 0.892774 |
| Height | 0.904 | 0.546975 | 0.983274 | 0.968536 |

TEM = Technical Error of Measurement, rTEM = relative TEM, R = Reliability coefficient, ICC = Intra class correlation coefficient

As shown in **Table 2.** the cross-validation results show a strong and statistically significant correlation between actual and estimated height in both data subsets, with a Pearson correlation of 0.865 in the 20% validation set and 0.792 in the 80% training set (both *p* < 0.001). The Mean Absolute Error (MAE) was slightly higher in the validation set (11.37 cm) compared to the training set (8.74 cm), indicating a modest increase in average prediction error when applied to unseen data. Similarly, the Standard Error of Estimate (SEE) was 14.30 cm in the validation set and 10.85 cm in the training set, reflecting greater variability in the residuals during validation. Despite these differences, the relatively small increase in error and strong correlations suggest that the model performs reliably and generalizes well across data partitions.

**Table 2:**
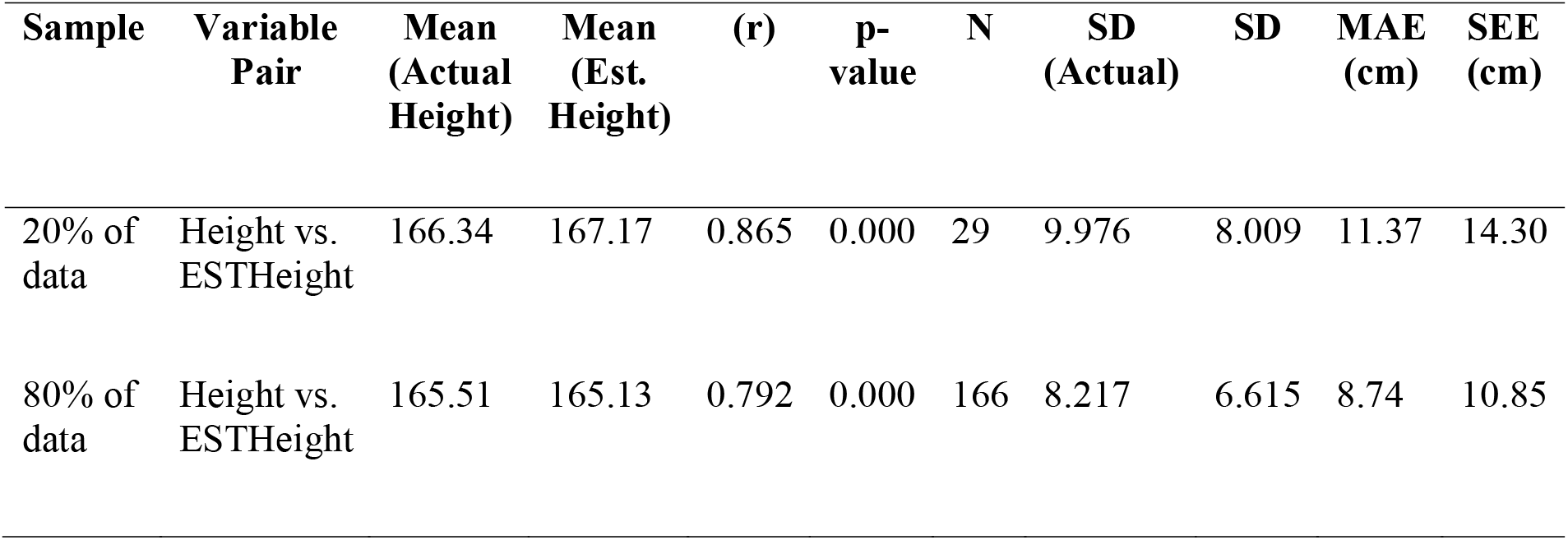
Cross validation between actual height and estimated height to display generalizability of data.

| Sample | Variable Pair | Mean (Actual Height) | Mean (Est. Height) | (r) | p-value | N | SD (Actual) | SD | MAE (cm) | SEE (cm) |
| --- | --- | --- | --- | --- | --- | --- | --- | --- | --- | --- |
| 20% of data | Height vs. ESTHeight | 166.34 | 167.17 | 0.865 | 0.000 | 29 | 9.976 | 8.009 | 11.37 | 14.30 |
| 80% of data | Height vs. ESTHeight | 165.51 | 165.13 | 0.792 | 0.000 | 166 | 8.217 | 6.615 | 8.74 | 10.85 |

### 2.4 Statistical Analysis

The IBM SPSS version 26 was used for data analysis. For descriptive statistics, the mean, standard deviation, maximum, minimum and standard error of the mean of the height, hand length and hand breadth were calculated for both the male and female populations. A paired t test was done to see if there was any significant difference between hand length and hand breadth of the right and left sides. Simple linear regression equations were derived from individual parameters, and R, R^2^ and Standard Error of Mean (SEE) were defined. A multiple linear regression equation was derived, including the hand length and hand breadth of both sides for both sexes, and it was used to calculate the estimated height. Spearman’s rho correlation coefficient was calculated to see the correlation between the actual height and estimated height.

## 3. Results

In this study 195 undergraduate medical students were enrolled; among them, 80 (41%) were female and 115 (59%) were male, with ages ranging from 18 to 24 years, with a mean age of 20.2 ± 1.35 years.

Table 3. Shows descriptive statistics of stature according to sex.

**Table 3:** Distribution of stature according to sex.

| Stature(cm) | Female (n = 80) | Male (n = 115) |
| --- | --- | --- |
| Mean | 159.01 | 170.24 |
| Std. Deviation | 7.261 | 5.794 |
| Std. Error of Mean | 0.812 | 0.540 |
| Minimum | 142 | 155 |
| Maximum | 188 | 188 |

Table 4. Illustrates descriptive statistics of hand length according to sex.

**Table 4:** Distribution of hand length according to sex.

| Hand length (cm) | Female |  | Male |  |
| --- | --- | --- | --- | --- |
|  | Right side | Left side | Right side | Left side |
| Minimum | 14.8 | 15.0 | 16.0 | 16.0 |
| Maximum | 19.4 | 19.5 | 21.5 | 21.5 |
| Mean | 16.861 | 16.897 | 18.482 | 18.570 |
| Std. Deviation | .9659 | .9487 | 1.0082 | 1.015 |

**Table 5**. Illustrates descriptive statistics of hand breadth according to sex.

**Table 5:** Distribution of hand breadth according to sex.

| Hand Breadth (cm) | Female |  | Male |  |
| --- | --- | --- | --- | --- |
|  | Right side | Left side | Right side | Left side |
| **Minimum** | 6.2 | 6.0 | 7.5 | 6.7 |
| **Maximum** | 9.1 | 9.2 | 9.9 | 9.7 |
| **Mean** | 7.311 | 7.229 | 8.356 | 8.264 |
| **Std. Deviation** | .5846 | .5818 | .4780 | .5034 |

Table 6. displays the outcome of a paired t-test conducted to see if RHL and LHL and, RHB and LHB differ significantly. A substantial positive association was discovered for pairs RHL and LHL (r = 0.984) and RHB and LHB (r = 0.953). that was statistically significant (p < 0.001). Hence, it was deduced that there was no discernible variation between the dimensions of the right and left hands that would have impacted the accuracy of the regression.

**Table 6:**
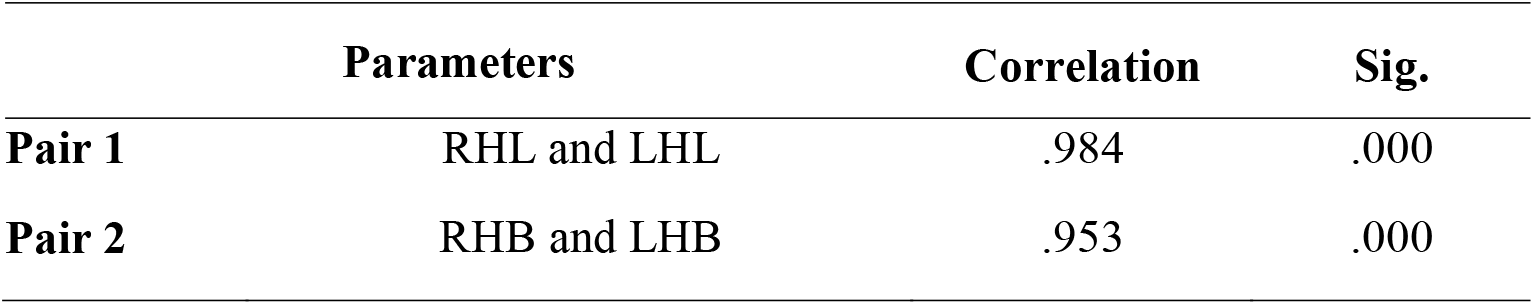
Paired t-test to see the differences between right-and left-hand dimensions.

For the estimation of stature from the length/breadth of the hand linear regression formula would be

Height (y) = a + b × Hand length/breadth (x)

[a = Constant and b = Independent variable]

**Table 7.** shows the linear regression equations derived for each parameter for males and females, along with their R, R^2^, SEE and p values.

**Table 7:** Linear regression equation for estimation of height from hand length and hand breadth in male and female.

| **Male** | **Female** |
| --- | --- |
| Height = $100.029 + (3.799 \times \text{RHL}^a)$ | Height = $74.114 + (5.035 \times \text{RHL}^a)$ |
| [R = .661, R<sup>2</sup> = .437, SEE = 4.4, p < 0.001] | [R = .670, R<sup>2</sup> = .449, SEE = 5.4, p < 0.001] |
| Height = $99.892 + (3.788 \times \text{LHL}^b)$ | Height = $73.668 + (5.051 \times \text{LHL}^b)$ |
| [R = .664, R<sup>2</sup> = .441, SEE = 4.4, p < 0.001] | [R = .660, R<sup>2</sup> = .435, SEE = 5.4, p < 0.001] |
| Height = $130.672 + (4.736 \times \text{RHB}^c)$ | Height = $145.720 + (1.818 \times \text{RHB}^c)$ |
| [R = .391, R<sup>2</sup> = .153, SEE = 5.4, p < 0.001] | [R = .146, R<sup>2</sup> = .021, SEE = 7.2, p = 0.195] |
| Height = $133.334 + (4.466 \times \text{LHB}^d)$ | Height = $139.505 + (2.699 \times \text{LHB}^d)$ |
| [R = .388, R<sup>2</sup> = .151, SEE = 5.4, p < 0.001] | [R = .216, R<sup>2</sup> = .047, SEE = 7.1, p = 0.054] |

<sup>a</sup>RHL = Right hand length, <sup>b</sup>LHL = Left hand length, <sup>c</sup>RHB = Right hand breadth, <sup>d</sup>LHB = Left hand breadth.

Regression equation for estimation of stature with hand length and hand breadth combined for males and female is as follows:

- Female: Height = 77.633 – (6.307 × RHB) + (4.857 × LHB) + (3.591 × RHL) + (1.884 × LHL)
- Male: Height = 94.462 + (1.254 × RHL) + (2.313 × LHL) + (1.065 × RHB) + (0.092 × LHB)

Table 8. explains the model summary of the regression equation analysis based on combined hand length and hand breadth of the males and females. It indicates a R of .702 for females; implying there is a moderate to strong correlation between hand dimensions and stature. There was 49.2% variance in female height, while the predictions deviate by 5.3cm on average. The R value for males was .671, which means there is moderate correlation between hand dimensions and stature. The model also displays that there is 45.1% variance in male height, and the predictions are about 4.4 cm off on average.

**Table 8:** Model summary obtained after regression analysis of hand length and hand breadth combined.

| Sex | Model | R | R Square | Adjusted R Square | Std. Error of the Estimate |
| --- | --- | --- | --- | --- | --- |
| Female | 1 | .702 | .492 | .465 | 5.310 |
| Male | 1 | .671 | .451 | .431 | 4.372 |

Multiple linear regression equation combining female and male is as follows:

Height (cm) = 69.652 – (1.084 × RHB) + (2.307 × LHB) + (2.394 × RHL) + (2.451 × LHL) [R = .807, R^2^ = .652, SEE = 5.056]

Table 9. shows the descriptive statistics of actual height and the estimated height derived from the multiple linear regression equation.

**Table 9:**
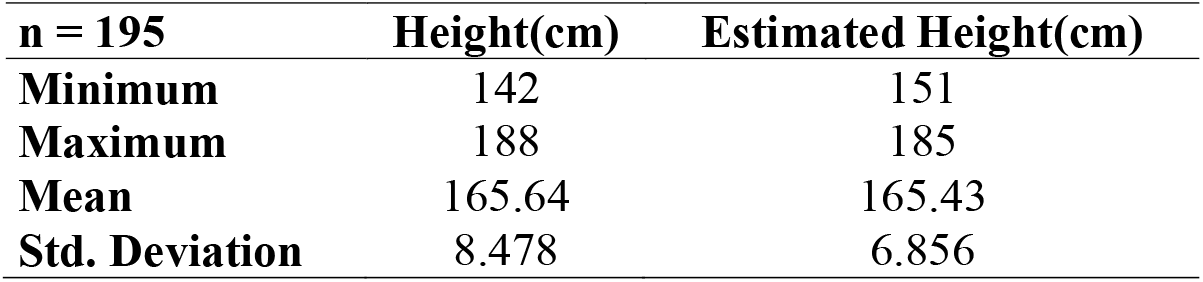
Descriptive statistics of actual height and estimated height.

| n = 195 | Height(cm) | Estimated Height(cm) |
| --- | --- | --- |
| Minimum | 142 | 151 |
| Maximum | 188 | 185 |
| Mean | 165.64 | 165.43 |
| Std. Deviation | 8.478 | 6.856 |

The Kolmogorov-Smirnov test was done, which deduced that data for actual height was not normally distributed (p < 0.001) but normally distributed (p = 0.76) for estimated heights, so that Spearman’s rho correlation was done. **Table 10**. displays the results. The correlation coefficient was 0.810, which signifies a strong positive correlation, and the values were statistically significant (p < 0.001).

**Table 10:** Spearman’s rho correlation between actual height and estimated height.

|  |  | Height (cm) | Estimated Stature (cm) |
| --- | --- | --- | --- |
| Height (cm) | Correlation Coefficient | 1.000 | .810 |
|  | Sig. (2-tailed) | . | .000 |
| Estimated Stature (cm) | Correlation Coefficient | .810 | 1.000 |
|  | Sig. (2-tailed) | .000 | . |

**Fig. 2.**
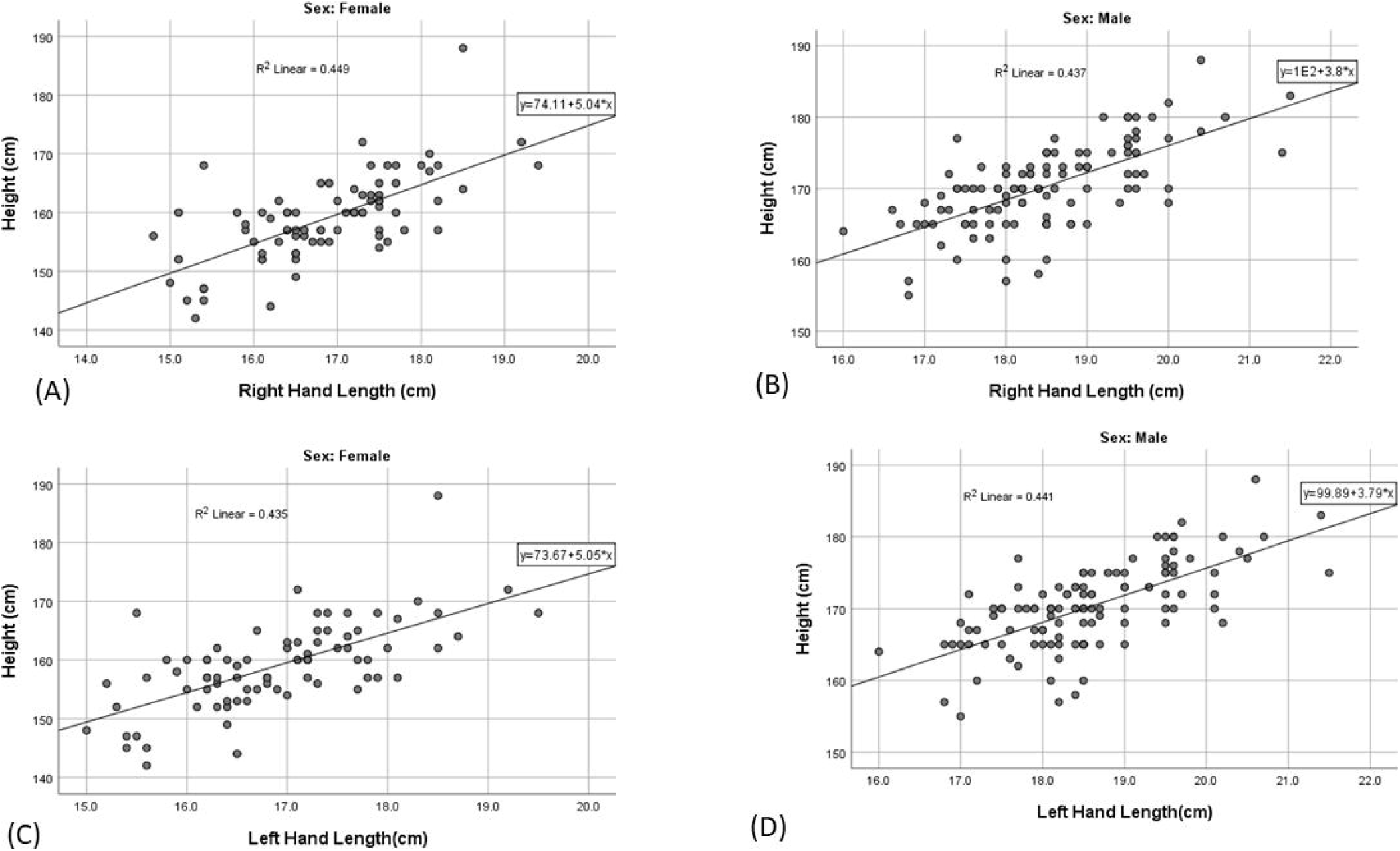
Scatter plot showing relationship between stature and hand lengths in male and female. (CI = 95%) (A) Between stature and RHL in females. (B) Between stature and RHL in males. (C) Between stature and LHL in females. (D) Between stature and LHL in males.

**Fig. 3.**
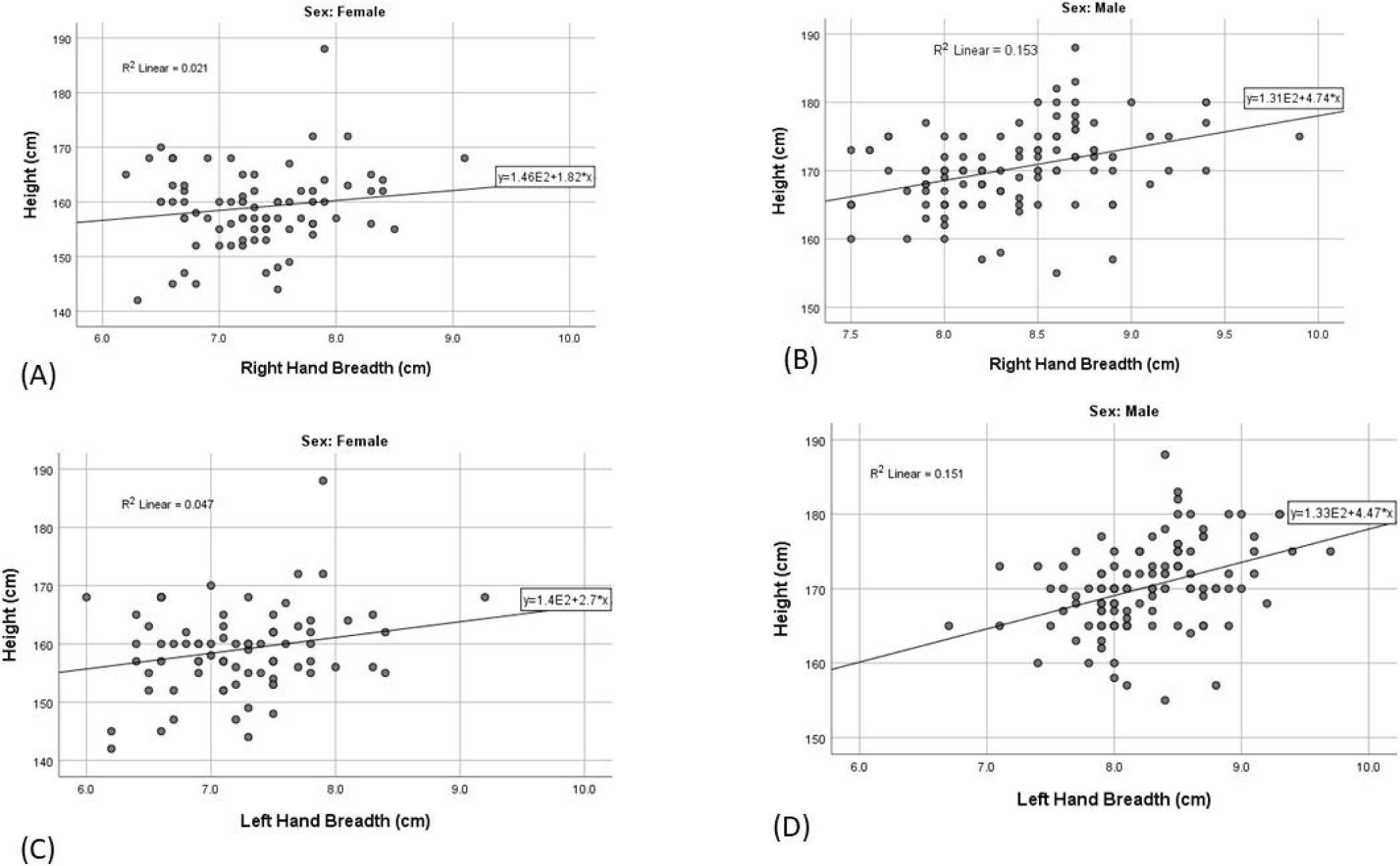
Scatter plot showing relation between stature and hand breadths in male and female. (CI = 95%) (A) Between stature and RHB in females. (B) Between stature and RHB in males. (C)Between stature and LHB in females. (D) Between stature and LHL in males.

## 4. Discussion

Stature is a changing value, and it depends upon various factors. This study attempted to determine if estimation of stature can be done solely by the length and breadth of the right and left hands. Mean length and breadth of both right and left hands were found to be higher in males (RHL = 18.48±1.00 cm, LHL = 18.57±1.01 cm, RHB = 8.36±4.78 cm and LHB = 8.26±5.03 cm) as compared with females (RHL = 16.861±0.96 cm, LHL = 16.897±0.95 cm, RHB = 7.31±0.58 cm, and LHB = 7.23±0.58 cm). Mean stature was also higher in males (170.24±5.79 cm) when compared to females (159.01±7.261 cm). It is similar to many other studies. [3–6,15–20] In a study done by Shakya et al., upper arm length of the participants was taken for stature estimation instead of hand length, and again, measurements were higher in males (Right = 41.4667±2.993 cm, Left = 40.598±2.849 cm) compared to females (Right = 37.436±1.807 cm, Left = 36.920±1.735 cm).[18] Male being genetically taller than female is the reason for this discrepancy.

Among the single-variable linear equations where stature was taken as the dependent variable, RHL and LHL both displayed the lowest SEE (4.4 cm) in males, whereas RHB and LHB, both had the same SEE (5.4 cm), and the results were statistically significant (p<0.001). Thus, hand lengths can be considered a more accurate predictor of stature in males compared to hand breadth. Similarly, in females, RHL and LHL had the lowest SEE (5.4 cm) and were statistically significant (p<0.001), whereas RHB had the highest SEE and was not statistically significant (p = 0.195). Thus, it can be deduced that RHL and LHL can be strong, accurate predictors of height in the female population. SEE was lower when all the variables were applied as explanatory variables in the case of males (4.37 cm) and females (5.3 cm) concluding that equation obtained from multiple parameters is more accurate. SEE in this study ranged from 4.4 to 7.2 cm; however, in a study done by Ozaslan in the Turkish population, SEE ranged from 6.03 to 6.39 cm [21] and much lower, 3.49 to 4.28 cm, in a study done by Pal et al.[7] In another study done by Habib and Kamal, in the Egyptian population, hand length and phalanges were used for stature estimation, SEE ranged from 5.3 to 7.27 cm, displaying values much similar to this study.[22] Rastogi et al. did a study in the Indian population and derived SEE ranging from 4.11 to 5.99 cm, which is also lower than the present study.[23] Sanli et al. did a study in the Turkish population using hand length as a variable for stature estimation and reported an SEE of 3.49 cm.[12] Panjokh et al. reported SEE of 4.87 in males and 5.11 cm in females in a multiple regression equation where the variables were forearm length, hand length and hand width these values align with this study.[19] In a study done by Shrestha et al., hand breadth showed a weak positive correlation, and hand length showed a moderate to strong positive correlation with stature, and they reported an SEE of 5.43 cm for males with an R^2^ of 0.125 and an SEE of 6.82 and R^2^ of 0.039 for the regression equation obtained from hand breadth and height[16] The values of R^2^ were much less for males than seen in this study (R^2^ = 0.153 for RHB and 0.151 for LHB) but almost similar for females (R^2^ = 0.021 for RHB and 0.047 for LHB).

As seen in the model summary of the multiple linear regression, R is .807, which indicates that there is a strong positive correlation between stature and hand dimensions. SEE is 5.056, indicating that the prediction deviates from the actual height by 5 cm on average. R^2^ is 0.652, and adjusted R^2^ is 0.644, depicting the robustness of the regression with p < 0.001 confirming that it is statistically significant. The correlation coefficient was higher in comparison to the study done by Mansur et al. (r = .703), in which a regression equation was derived using foot length[14] In another study done by Shrestha et al. craniometric analysis was used to derive a multivariate regression equation with r = 0.590.[24] In addition, another study derived a regression equation using percutaneous tibial length, and the correlation coefficient was found to be 0.62 for males and 0.71 for females.[25] Thus, it can be deduced that hand length and hand breadth could be a strong predictor of stature compared to other body parts. The linear regression model using hand length and hand breadth or hand length alone was found to be valid in similar previous studies.[4,15] Pandeya et al. took only hand length as a parameter for their estimation of the stature with R^2^ of 0.372 for male, 0.556 for female and 0.250 combined[15] Comparing these values with the present study it can be deduced that the multiple regression equation derived from this study is a stronger predictor of height. This study included hand breadth as a parameter that might have led to this significance. The R and R^2^ and the SEE values of the multiple linear regression equation were almost similar to the values obtained from equations of single variables, except for right hand breadth and left hand breadth in females, where R and R^2^ were very low and SEE was higher compared to equations obtained from other single variables. Pal et al. deduced that a multiple linear regression equation where explanatory variables were hand length, palm length, hand breadth and maximum hand breadth, with stature as the dependent variable, had higher reliability with lower SEE and high R and R^2^ values when compared to the equations obtained from single variables.[7] Palm length and maximum hand breadth were two extra variables included in Pal’s study compared to this study that may have caused this difference in the multiple and single variable linear regression equations. Thus, it can be deduced that using multiple parameters to generate a multiple regression equation can be a stronger predictor of height when compared to a single parameter.

Spearman’s rho correlation coefficient between height and estimated height was 0.810 (p < 0.001), confirming that with the increase in the measured height, estimated height also increased. Thus, it was deduced that the estimated height obtained from the multiple regression equation can be a proxy for the actual height.

In future, similar studies can be done using participants from various age groups so that age factors can also be analysed in the estimation of stature. The hand length and breadth were used as variables in this study, while some studies show that SEE can be decreased using more variables like foot length and forearm and arm length, etc. This study paves a way for such studies to be done in future.

Summary of regression analysis of various studies is shown in **Table 11**. It can be compared to present study referring to **Table *7***.

**Table 11:**
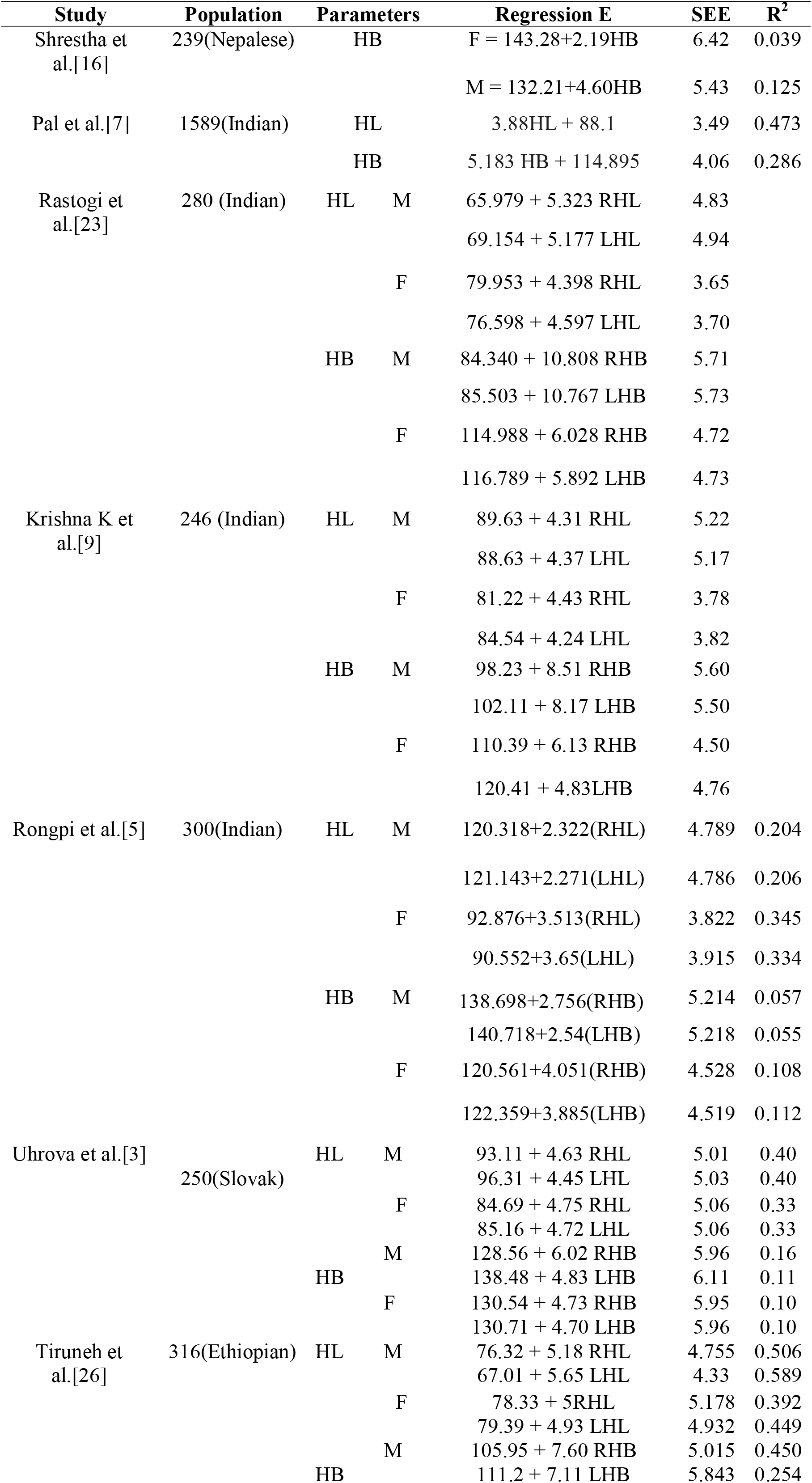
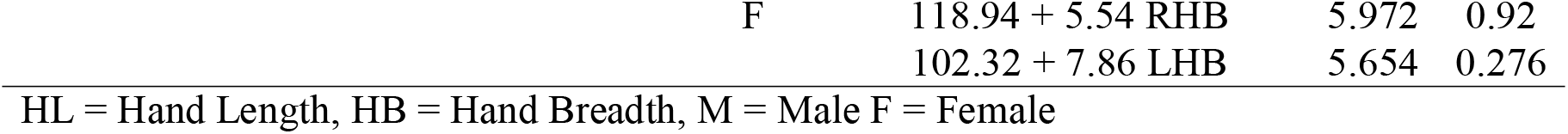
Summary of regression analysis of previous studies in comparison to this study.

## 6. Limitations

This study was done on a limited number of students at a medical college, so in order to decrease bias and add more significance, a larger population sample can be taken in the future. It included individuals of age groups 18 – 24 and cannot represent the entire population of various age groups. This is acknowledged as another limitation of this study.

## 6. Conclusions

Stature is an important entity for personal identification of the individual. Various studies have been done in the past to derive equations for estimation of stature from different body parts. This study was done on medical students, and it generated regression equations for estimation of stature by taking into account hand length and hand breadth. Mean length and breadth of hand and stature were seen to be more in males than in females. The multiple linear regression equation unveiled that there is a strong positive correlation between stature and the hand dimensions. Similar studies can be done in the future, taking a larger population size and participants from different age groups and also considering parameters from different body parts.

## 7. Abbreviations

RHL: Right Hand Length
LHL: Left Hand Length
RHB: Right Hand Breadth
LHB: Left Hand Breadth
SEE: Standard Error of Estimate

## Acknowledgement

We would like to acknowledge medical students of Devdaha Medical College and Research Institute, Devdaha, Rupandehi, Nepal, who consented for participation in the study and Mr. Keshav Rajbhandari from Madan Bhandari Academy of Health Sciences, for helping us with the statistical analysis.

## Notes

Declaration of conflicting interest: The authors declare no potential conflicts of interest with respect to the research, authorship, and/or publication of this article.

### Competing Interest Statement

The authors have declared no competing interest.

### Summary of Updates

Abstract has be updated and the date of the conduction of research has been changed Method section has been updated and data reliability portion has been addded.

